# Composition of the holdfast polysaccharide from *Caulobacter crescentus*

**DOI:** 10.1101/602995

**Authors:** David M. Hershey, Sara Porfírio, Bernhard Jaehrig, Christian Heiss, Parastoo Azadi, Aretha Fiebig, Sean Crosson

## Abstract

Surface colonization is central to the lifestyles of many bacteria. Exploiting surface niches requires sophisticated systems for sensing and attaching to solid materials. *Caulobacter crescentus* synthesizes a polysaccharide-based adhesin known as the holdfast at one of its cell poles, which enables tight attachment to exogenous surfaces. The genes required for holdfast biosynthesis have been analyzed in detail, but an inability to isolate sufficient quantities of holdfast has limited efforts to characterize its composition and structure. In this report we describe a method to extract the holdfast from *C. crescentus* cultures and present a survey of its carbohydrate content. Glucose, 3-*O*-methylglucose, mannose, N-acetylglucosamine and xylose were detected in our extracts. Our results suggest that the holdfast contains a 1,4-linked backbone of glucose, mannose, N-acetylglucosamine and xylose that is decorated with branches at the C-6 positions of glucose and mannose. By defining the monosaccharide components in the polysaccharide, our work establishes a framework for characterizing enzymes in the holdfast pathway and provides a broader understanding of how polysaccharide adhesins are built.

## Introduction

Bacteria routinely encounter solid objects in their surrounding environments that present attractive niches for colonization (1, 2). Attachment to and growth in association with these surfaces is fundamental to survival for many bacteria (3). The diversity of potential colonization substrates ranges from small soil particles, to plant and animal tissues, to the interior walls of pipes and tanks in industrial settings (4). As such, surface associated bacteria present an important obstacle to many clinical, agricultural and manufacturing processes. Understanding how attached communities form and proliferate has the potential to inform strategies for manipulating surface colonization in order to increase economic efficiency.

Surface colonization is a stepwise process (5). Simply contacting a surface requires bacteria to overcome repulsive forces within the hydrodynamic layer that forms at the liquid-solid interface (6). In many cases, active processes such as flagellar motility and pilus oscillation provide the mechanical force needed to achieve direct contact (7-10). Transient interactions between various features displayed on the cell surface serve to stabilize this initial association, allowing for the deployment of dedicated adhesins that promote permanent attachment (11, 12). Bacterial adhesins are extremely diverse, displaying a range of chemical, functional and mechanical properties (13). Yet, irreversible attachment mediated by adhesins is thought to represent the committed step in surface colonization for many organisms (14, 15). It provides cells with the opportunity to grow and divide in association with the target substrate, which can lead to the formation of biofilms.

In the class *Alphaproteobacteria,* adhesins are often localized to one cell pole, giving the cell a defined geometry for attachment with respect to the surface substrate (16-20). *Caulobacter crescentus* is a dimorphic organism that has become a model for polar adhesion in this clade (13). During the *C. crescentus* cell cycle, sessile stalked cells divide asymmetrically to release a motile daughter called a swarmer cell that has pili and a flagellum at one pole (21). Swarmer cells remain motile for a period of time before shedding their pili and flagella and transitioning to stalked cells (22). At developmentally controlled times during the swarmer to stalked transition or in response to a physical encounter with a surface, *C. crescentus* can produce a specialized adhesin called the holdfast at the old cell pole (Fig 1A) (23-25). As the swarmer cell transitions, the holdfast remains displayed at the tip of the newly formed stalk where it is primed for surface attachment (24). The holdfast is one of the strongest adhesives characterized to date (26). It is gelatinous in nature, has elastic characteristics and can adhere to substrates that have a range of physical and chemical properties, making it an attractive model for developing adhesives to be used in medical and industrial settings (27, 28).

**Figure 1.**
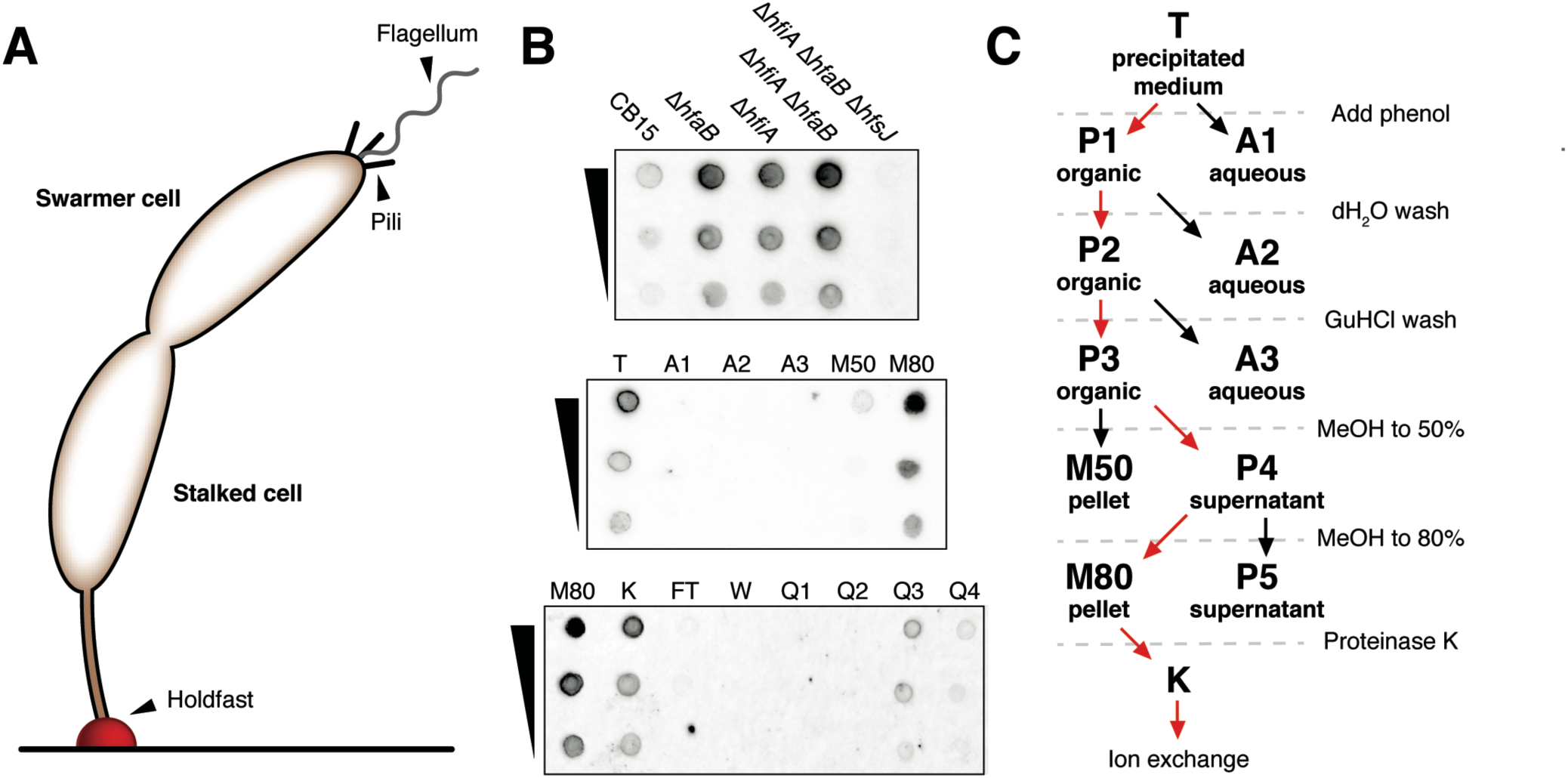
*Extraction of the* C. crescentus *holdfast*. (A) A surface attached *C. crescentus* cell at the pre-divisional stage. The holdfast adhesin (red) is depicted at the tip of the stalked cell, attached to a horizontal surface. Stalked cells divide to release a motile swarmer cell that possesses a flagellum and type IV pili. The released swarmer cell eventually transitions into a stalked cell and secretes a holdfast. (B) Enrichment of the holdfast polysaccharide from *C. crescentus* culture medium. Top: Lectin (fWGA)-stained dotblot measuring holdfast released into the spent medium by various mutants grown in PYE. Holdfast was extracted from spent medium by 60% ethanol precipitation, re-suspended in TES buffer (Materials and Methods) and spotted on nitrocellulose. Middle: Phenol extraction of precipitated medium from the Δ*hfiA* Δ*hfaB* strain. The re-suspended ethanol precipitation (T) was extracted with phenol. The aqueous phases (A samples) and subsequent methanol precipitates (M samples) are shown. Bottom: The M80 fraction was treated with proteinase K (K sample) and subjected to anion exchange chromatography. The flow-through (FT), wash (W) and step gradient fractions (Q1, Q2, Q3 and Q4) are shown. (C) Schematic of the holdfast enrichment procedure. Fraction names correspond to those shown in panel B.

Despite significant advances in understanding mechanical properties of the holdfast, its chemical composition remains poorly characterized. The adhesin binds to the N-acetylglucosamine (GlcNAc) specific lectin wheat germ agglutinin (WGA) and is sensitive to the GlcNAc specific hydrolases lysozyme and chitinase, indicating that it contains GlcNAc (16). Treatment with proteinase K or DNAse also alters the mechanical properties suggesting that protein and DNA may be included in the matrix as well (29). Finally, the genetic determinants of holdfast biosynthesis provide invaluable insight into its chemical makeup. Genes predicted to encode machinery for the production of an extracellular polysaccharide represent the major factors required for holdfast production (30). Among these genes are four predicted glycosyltransferases (GTs), implying that a complex polysaccharide with a repeating unit of at least four monosaccharides is a critical component of the holdfast matrix (31-33).

Extracellular polysaccharides are deployed by bacteria for a variety of purposes, including both as adhesins and as components of biofilm matrices (34). The polar polysaccharide adhesins secreted by *Alphaproteobacteria* are unusual both for their discrete localization within the cell envelope and their exceptional adhesive capabilities (13). Understanding the molecular basis for interactions between polar adhesins and exogenous surfaces requires information about the chemical structure of the adhesive matrix. However, both the insoluble nature of these materials and the limited amounts secreted by producing organisms have hindered efforts to obtain suitable samples for analysis. Here we describe a method to extract the holdfast from *C. crescentus* cultures. By analyzing the carbohydrate content of this preparation, we have identified the monosaccharide components of the holdfast and performed a preliminary characterization of their linkage patterns. The results are consistent with a model in which a 1,4-linked backbone of glucose, GlcNAc, mannose and xylose that is decorated with branches at the C-6 positions of glucose and mannose makes up the holdfast polysaccharide.

## Results

### Biochemical enrichment of the holdfast

*C. crescentus* cells produce only a small patch of holdfast that is tightly associated with the cell envelope, presenting a major challenge to isolating sufficient quantities of material. We took a genetic approach to engineer a strain that would simplify the extraction process. We focused on the *holdfast inhibitor A* (*hfiA*) gene, which limits the number of adhesive cells by inhibiting holdfast biosynthesis, and the *hfaB* gene that is required for anchoring the holdfast matrix to the cell pole (32, 35, 36). We reasoned that Δ*hfiA* Δ*hfaB* cells should overproduce holdfast and release it into the spent medium, allowing for the separation of holdfast from cell material by centrifugation. To assess this strategy, the spent medium from a set of *C. crescentus* strains was precipitated with ethanol and probed by dotblotting with fluorescently-labeled WGA (fWGA) (Fig 1B). Wild-type *C. crescentus* released a small amount of WGA reactive material, and the signal from Δ*hfaB* cultures was higher, consistent with the previously reported shedding phenotype in this mutant (36). Surprisingly, the Δ*hfiA* mutant also released a significant amount of WGA reactive material, suggesting that hyper-holdfast production in this background results in the release of holdfast from the cell surface even in the presence of an intact anchor complex. Finally, the Δ*hfiA* Δ*hfaB* double mutant released more WGA reactive material than the Δ*hfiA* or Δ*hfaB* single mutants, and the signal was eliminated when the *hfsJ* gene, encoding a GT required for holdfast biosynthesis, was deleted in the Δ*hfiA* Δ*hfaB* background (32).

We precipitated the spent medium from Δ*hfiA* Δ*hfaB* cultures that had been grown in defined M2X medium to minimize contaminants from the medium. The precipitated WGA-reactive signal migrated to the organic phase of a phenol extraction and could be precipitated from the phenol phase by the addition of increasing amounts of methanol. To further enrich the holdfast polysaccharide, we first treated precipitated phenol extracts with proteinase K. WGA staining as detected by dotblot decreased by ∼50% upon protease treatment. Unknown protein(s) have been shown to contribute to the mechanical properties of the holdfast, and the digestion of these proteins may alter either how the material binds to the nitrocellulose membrane or the accessibility of GlcNAc residues within the matrix to WGA (29). The protease digests were subsequently subjected to anion exchange chromatography. The WGA-reactive signal bound to Q sepharose resin and could be eluted with solutions of increasing ionic strength. The peak fraction was dialyzed extensively against H_2_O and used for subsequent analysis (Fig 1C).

The UV absorbance spectrum of a typical holdfast preparation is shown in Fig 2. No peaks at 260nm or 280nm were present demonstrating that the preparation is free of significant contamination from protein and nucleic acids. We also examined the extract using light microscopy. Though no phase contrast signal was detected in the sample, discrete foci were observed when fWGA was added, reminiscent of the staining signal from cell-associated holdfasts (Fig 3). To confirm that the fluorescent foci were derived from holdfast, we stained an equivalent extract prepared from the Δ*hfiA* Δ*hfaB* Δ*hfsJ* strain. Because this sample was not expected to have significant signal in the fWGA channel, fluorophore-conjugated beads were added in order to define the focal plane. When viewed with a GFP filter, the fluorescent beads were evenly dispersed across the slide in the Δ*hfiA* Δ*hfaB* Δ*hfsJ* extracts, but no fWGA foci were present in the red channel. We concluded that the extraction yielded discrete WGA-reactive particles that could be extracted from cells with an intact holdfast synthesis pathway.

**Figure 2.**
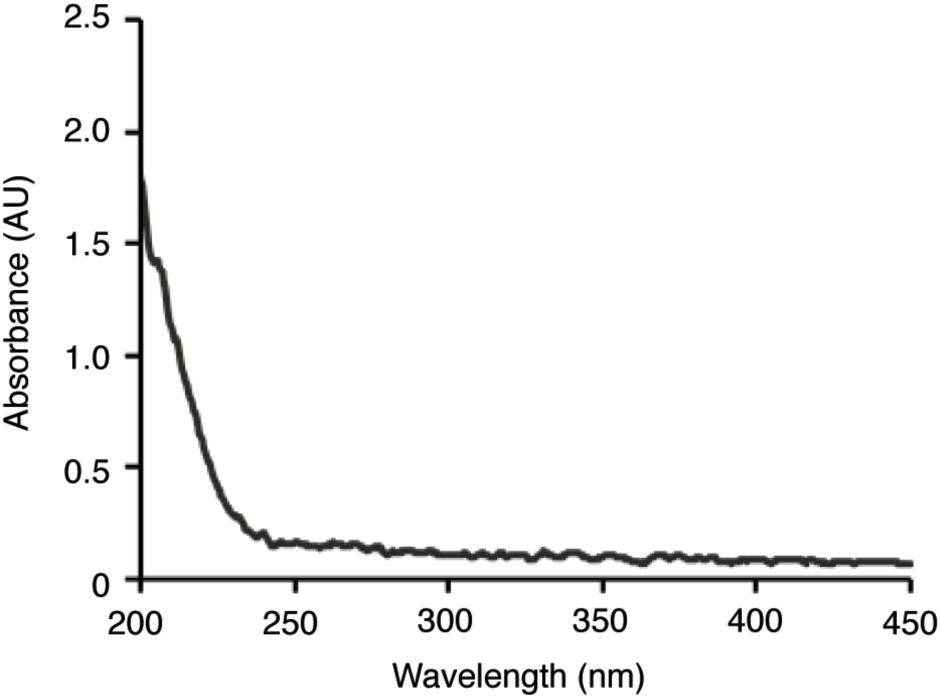
UV-vis absorbance spectrum of a typical holdfast extract. An absorbance spectrum lacking peaks in the 260nm or 280nm ranges demonstrated that the sample used for analysis was free of significant protein or nucleic acid contamination.

**Figure 3.**
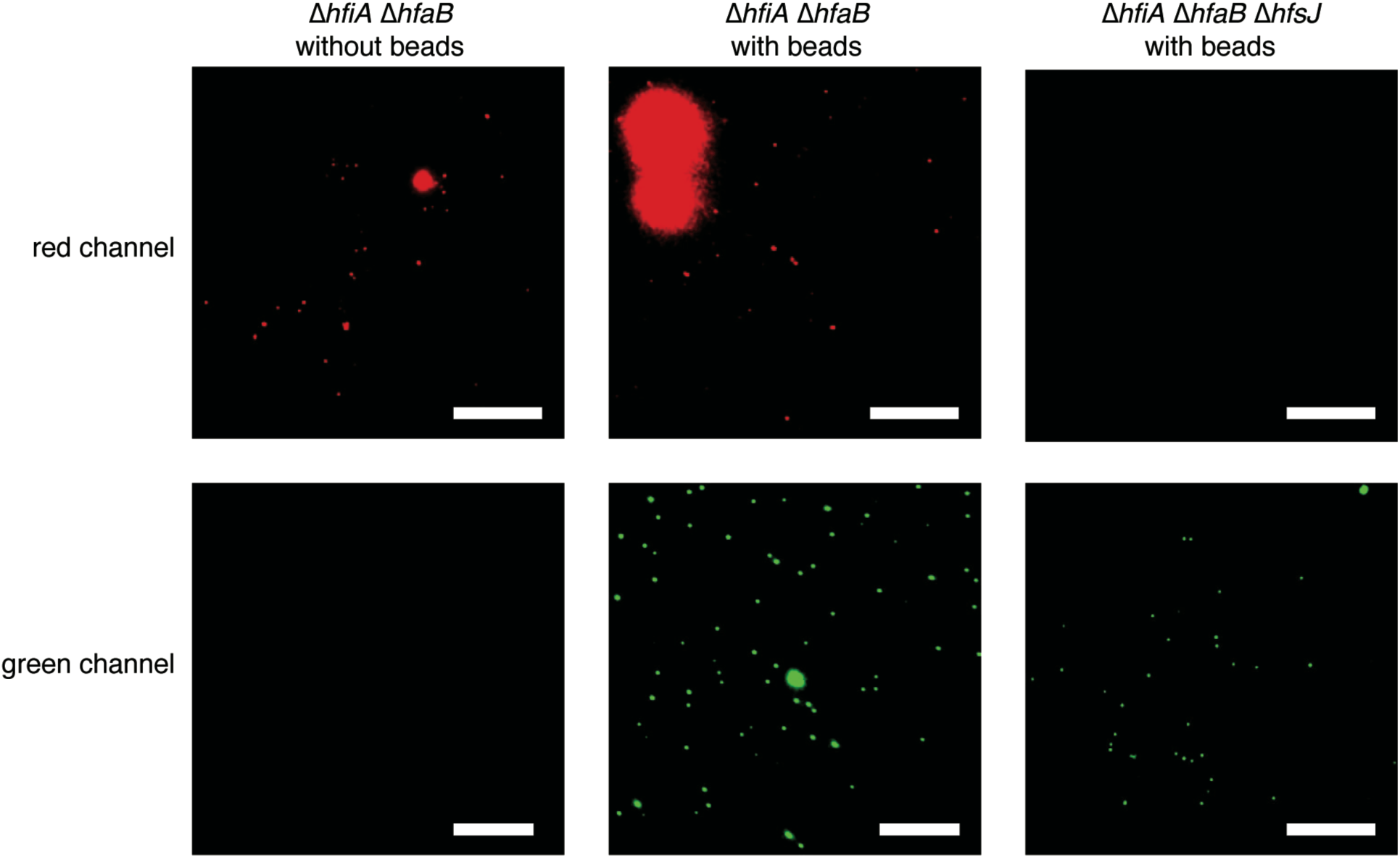
WGA-reactive particles in holdfast extracts. Extracts prepared from the indicated strains were stained with fWGA in the presence or absence of fluorescently labeled beads. Top panels: red channel (Alexa594-WGA); bottom panels: green channel (0.2µm YP microbeads). The scale bars represent 10µm.

### Monosaccharide composition of the holdfast

To examine the monosaccharide content of the holdfast we first prepared trimethylsilyl (TMS) methyl glycosides from acid-hydrolyzed extracts and analyzed them by gas chromatography-mass spectrometry (GC-MS). Because this method can identify neutral monosaccharides, amino sugars and uronic acids, its broad scope is ideal for exploratory analysis. The preliminary composition profile identified glucose, GlcNAc, mannose, xylose and a methylated hexose (data not shown). Guided by these results, we chose to prepare alditol acetates from acid-hydrolyzed holdfast extracts and analyze both neutral and amino sugar derivatives by GC-MS. Alditol acetate derivatization provides a more sensitive view of monosaccharide composition and can also be used to define the structure of methylated sugars. We confirmed that glucose, GlcNAc, mannose and xylose were present and also characterized the methyl-hexose as 3-*O*-methylglucose (Table 2). Yields from the holdfast extraction procedure were too low to weigh the material accurately, and we were unable to determine the total carbohydrate content in the samples.

**Table 1.**
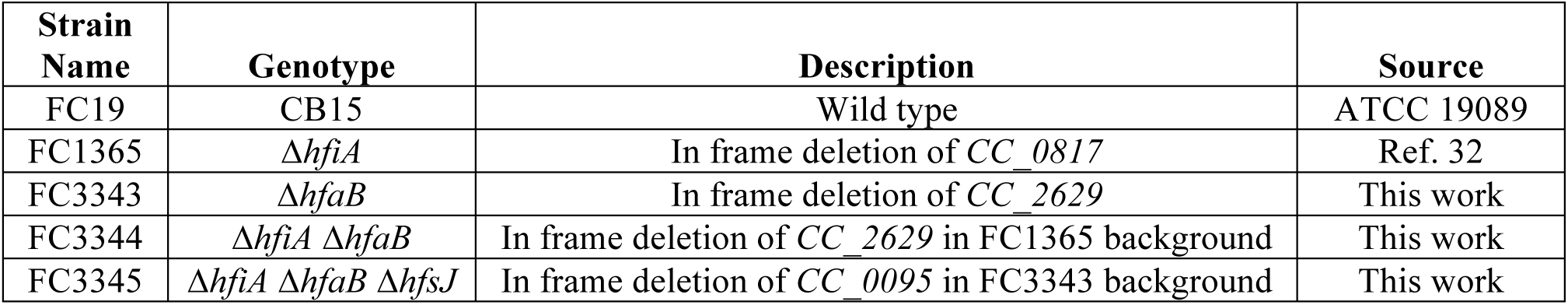
Caulobacter crescentus *strains used in this study*

| Strain Name | Genotype | Description | Source |
| --- | --- | --- | --- |
| FC19 | CB15 | Wild type | ATCC 19089 |
| FC1365 | $\Delta hfiA$ | In frame deletion of *CC_0817* | Ref. 32 |
| FC3343 | $\Delta hfaB$ | In frame deletion of *CC_2629* | This work |
| FC3344 | $\Delta hfiA \Delta hfaB$ | In frame deletion of *CC_2629* in FC1365 background | This work |
| FC3345 | $\Delta hfiA \Delta hfaB \Delta hfsJ$ | In frame deletion of *CC_0095* in FC3343 background | This work |

**Table 2.** Monosaccharide composition of a typical holdfast extract. Values are reported as mean ± standard error from two biological replicates.

| **Monosaccharide** | **Mole percent** |
| --- | --- |
| 3-*O*-methylglucose | 12 ± 3 |
| Glucose | 15 ± 4 |
| Mannose | 18 ± 1 |
| N-acetylglucosamine | 34 ± 4 |
| Xylose | 20 ± 7 |

The identification of xylose drew our attention because the cultures used for holdfast extractions had been grown in M2 salts with xylose as the carbon source. Though it seemed unlikely, we wanted to confirm that the xylose detected in our samples was not derived from the growth medium. We repeated the isolation procedure using Δ*hfiA* Δ*hfaB* cultures that had been grown in peptone-yeast extract (PYE), a complex medium containing mainly amino acids as the carbon source. The neutral sugar profiles of hydrolyzed holdfast from cells grown in M2X and PYE medium were nearly identical, and xylose was present in both samples (Fig 4). We conclude that the holdfast contains glucose, GlcNAc, mannose, xylose and 3-*O*-methylglucose and that the growth medium has little effect on this composition profile.

**Figure 4.**
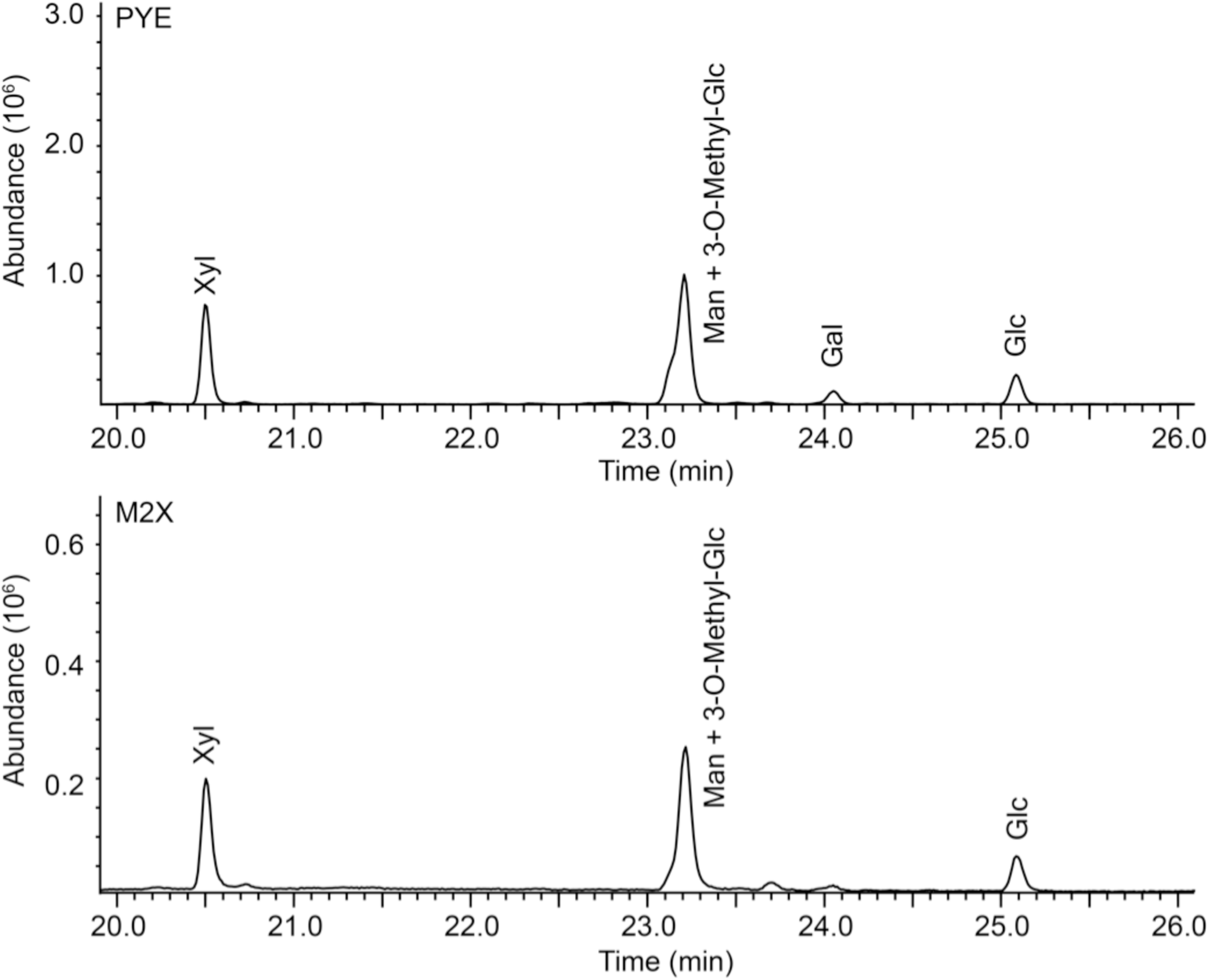
Comparison of neutral sugar profiles in holdfast extracts from two different growth media. Total ion chromatograms from GC-MS analysis of extracts from cell grown in PYE (top) or M2X (bottom) medium are shown. Xylose (Xyl) was present in both samples demonstrating that this component is not derived from xylose in the medium. The similarity in the monosaccharide profiles indicates minimal effects of medium type on holdfast composition. Mannose (Man); 3*-O*-methylglucose (3-O-Methyl-Glc); glucose (Glc); galactose (Gal).

### Determination of monosaccharide linkages

To characterize the linkage patterns of monosaccharides in the holdfast extract, we prepared partially methylated alditol acetates (PMAAs) and analyzed them by GC-MS. Because neutral monosaccharides and amino sugars cannot be separated efficiently with the same GC method, we analyzed the two sets of PMAAs on separate columns. We merged the two datasets by normalizing peak area percentages to the signal for 4-linked glucose, which was present in both chromatograms (Table 3). The analysis identified 4-linked forms of glucose, GlcNAc, mannose and xylose as the major residues. Terminal galactose, terminal glucose, terminal GlcNAc, terminal mannose, terminal xylose, 2-linked xylose, 4,6-linked mannose and 4,6-linked glucose were detected as minor components.

**Table 3.** Analysis of monosaccharide linkages from a typical holdfast extract using CH_3_I derivatization.

| **Residue** | **Normalized peak percent** |
| --- | --- |
| Terminal galactopyranosyl | 1 |
| Terminal glucopyranosyl | 13 |
| Terminal mannopyranosyl | 2 |
| Terminal N-acetylglucosaminopyranosyl | 2 |
| Terminal xylopyranosyl | 1 |
| 2-linked xylopyranosyl | 4 |
| 4-linked glucopyranosyl | 27 |
| 4-linked mannopyranosyl | 18 |
| 4-linked N-acetylglucosaminopyranosyl | 22 |
| 4-linked xylopyranosyl | 7 |
| 4,6-linked glucopyranosyl | ~1 |
| 4,6-linked mannopyranosyl | 1 |

To perform linkage analysis, the hydroxyls in an intact polysaccharide are methylated, the methylated polysaccharide is hydrolyzed, and the resulting partially methylated monosaccharides are reduced, acetylated and analyzed by GC-MS. Both glucose and 3-*O*-methylglucose are present in the holdfast, and methylation of the polysaccharide renders PMAAs derived from glucose and those derived from 3-*O*-methylglucose indistinguishable. To define the linkage state of 3-*O*-methylglucose, we repeated the linkage analysis by treating the holdfast polysaccharide with CD_3_I in instead of CH_3_I, which is normally used to prepare PMAAs. This allowed us to use mass spectrometry to differentiate methyl groups in the native polysaccharide from those added during derivatization. The neutral sugar profile obtained with CD_3_I methylation was similar to that obtained using CH_3_I (Tables 3 and 4). 4-linked glucose, 4-linked mannose and 4-linked xylose were the major components. Terminal 3-*O*-methylglucose, terminal xylose, 4,6-linked glucose and 4,6-linked mannose were minor components. We identified terminal 3-*O*-methylglucose by the presence of 121, 165, 167 and 211 (m/z) ions (Fig 5), demonstrating that the methyl groups in the holdfast polysaccharide are attached to C3 of terminal glucose residues.

**Table 4.** Analysis of neutral monosaccharide linkages from a typical holdfast extract using CD_3_I derivatization

| **Residue** | **Peak percent** |
| --- | --- |
| Terminal 3-*O*-methylglucopyranosyl | 9 |
| Terminal xylopyranosyl | 12 |
| 4-linked glucopyranosyl | 21 |
| 4-linked mannopyranosyl | 27 |
| 4-linked xylopyranosyl | 23 |
| 4,6-linked glucopyranosyl | 4 |
| 4,6-linked mannopyranosyl | 5 |

**Figure 5.**
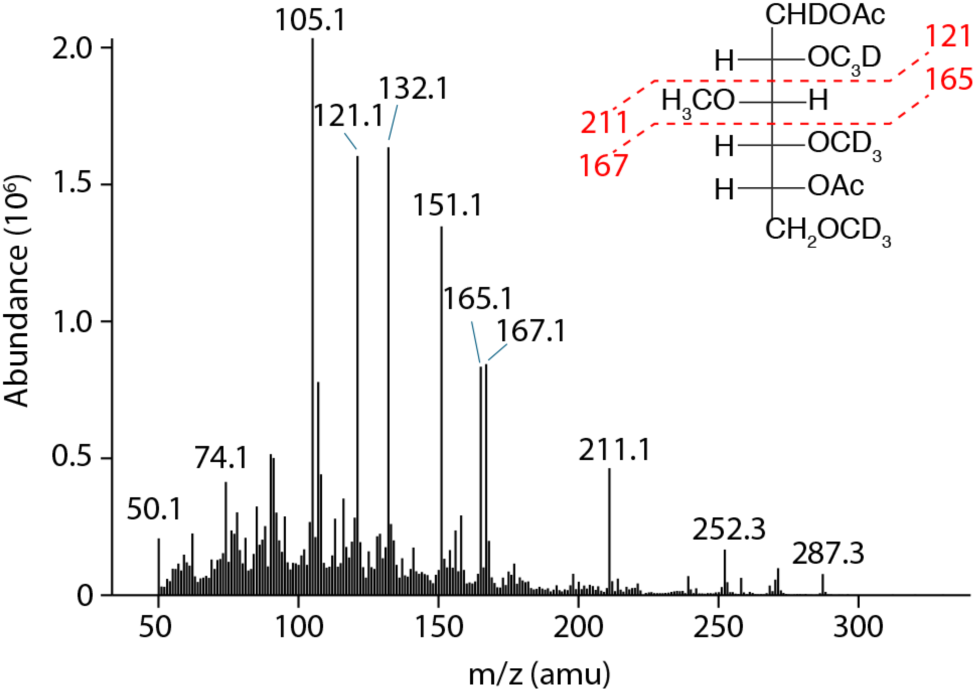
Mass spectrum of PMAA derived from terminal-3-O-methylglucopyranose. The predicted fragmentation pattern of a CD_3_I treated derivative of terminal 3-*O*-methylglucopyranose is shown in the top right corner. Characteristic ions used to define the position of the polysaccharide-derived methyl group are shown in red.

## Discussion

Permanent attachment is a critical stage in the process of surface colonization by bacteria (37). As the first irreversible step, it represents the point at which an organism commits to colonizing a substrate (38). A unipolar adhesin called the holdfast is the primary determinant of surface colonization in the model alphaproteobacterium *Caulobacter crescentus* (33). Biophysical studies of the *C. crescentus* holdfast have confirmed the material’s high versatility and adhesive strength, while also providing evidence for a layered substructure within the matrix (28, 29). Detailed genetic analysis of holdfast synthesis has shown that production of an extracellular polysaccharide is the key determinant (30). Binding of the GlcNAc specific lectin WGA to the holdfast matrix and its sensitivity to lysozyme and chitinase further support the notion that it is a polysaccharide-based material (16).

Linking the mechanical properties of the holdfast to specific chemical features has been limited by a striking lack of information about its composition. To understand the molecular basis for holdfast-based adhesion and to begin reconstituting the biosynthetic pathway, chemical analysis of the polysaccharide is needed. Despite decades of interest, such experiments have been precluded by an inability to isolate the material. We took a genetic approach to streamline the extraction process by deleting the genes for the holdfast inhibitor HfiA and a component of the holdfast anchor, HfaB. The resulting strain produced more holdfast than wild type and released it from the cell surface, allowing for the enrichment of holdfast material directly from spent medium (Fig 1). This approach yielded a sample largely free of protein and nucleic acid that could be used for carbohydrate analysis (Fig 2). More broadly, the extraction we describe here provides a framework for isolating unipolar adhesins that can be optimized and scaled to facilitate more detailed structural studies.

Glucose, GlcNAc, mannose, and xylose were the major monosaccharides in our preparations. These residues were predominantly 4-linked leading us to propose that they constitute a 1,4-linked backbone within the holdfast polysaccharide. Determining the order of residues in this chain will require more nuanced approaches including NMR analysis, which can be undertaken once sufficient material has been obtained. We also identified 3-*O*-methylglucose as a minor, terminally linked component of the holdfast. Methylation of glucose at the non-reducing end of the polysaccharide could serve as a termination signal for polymerization as it does for certain ABC transporter-dependent synthesis pathways (39, 40). However, the relative abundance of 3-*O*-methylglucose suggests that it is more likely a sub-stoichiometric component of the repeating unit and not present in a single copy exclusively at the chain terminus. We propose that 3-*O*-methylglucose sits at the termini of branches within the repeating unit. Indeed, we consistently detected 4,6-linked glucose and 4,6-linked mannose in our samples indicating that glucose and mannose residues within the 1,4-linked backbone are substituted at C6. The identification of glucose and mannose in both the 4- and 4,6-linked forms suggests that C-6 substitutions at these residues are non-stoichiometric and that branching is heterogeneous.

We observed some variability among the less abundant signals in our extracts. Glucose, GlcNAc, mannose and xylose, though primarily 4-linked, were also detected in terminally linked forms at low levels, and the presence of these signals was inconsistent (Tables 3 and 4). It seems unlikely that small amounts of each monosaccharide are present at branch termini. Instead these signals likely reflect non-specific hydrolysis of the 1,4-linked backbone during either the culture growth or extraction processes. Trace amounts of galactose were also detected only in some extracts, and we identified both 2-linked xylose and terminal galactose in samples containing galactose. This suggests that some branches may contain xylose substituted at C2 with galactose. Variability in the linkage analysis is likely due in part to the difficulty of quantitatively permethylating and hydrolyzing regions with branched residues, particularly with limited quantities of sample. Determining the exact nature of the C6 substitutions on glucose and mannose will require optimizing the extraction described here to increase yields.

The genes required for holdfast production encode a predicted *wzy*-type polysaccharide assembly pathway (30, 33). In the first stage of this process, GT enzymes add monosaccharides to a lipid carrier, producing a glycolipid repeating unit. Enzymatic machinery located in the cell envelope then polymerizes the repeating unit and secretes the resulting polysaccharide to the cell surface (41, 42). Though the polymerization and export machinery is conserved among bacteria, each pathway uses a characteristic set of GTs with varying specificities to assemble a repeating unit, explaining the enormous diversity of *wzy*-dependent carbohydrates (43-45). Extensive genetic analysis using saturating mutagenesis has shown that four GTs are required for holdfast production (31-33), but our results suggest that repeating units with up to six or even seven sugars may be present in the polysaccharide. GTs that perform multiple monosaccharide additions using a single active site have been described (46-48), and such activities may explain how four enzymes synthesize a repeating unit with greater than four sugars. For instance, a bifunctional GT might both elongate the 1,4-linked backbone and introduce a branch by sequentially adding monosaccharides to C-4 and C-6 of the terminal sugar in a glycolipid intermediate. Alternatively, GTs required for C-6 branching may not be essential for holdfast production. Mutating these genes could disrupt branching without affecting the adhesive properties of the holdfast, making them difficult to identify using forward genetics. *In vitro* studies with the holdfast GTs should help to clarify this point.

Though polysaccharide production is the main genetic determinant of holdfast-based adhesion, there is evidence that other macromolecules are associated with the matrix (29). Proteinase K treatment alters the elastic properties of surface-attached holdfasts, and our biochemical fractionation supports the presence of protein (29). Not only did treating holdfast extracts with proteinase K reduce their WGA reactivity, but the material also generally partitioned like a protein; it migrated to the organic phase of a phenol extraction and precipitated in phenol with high concentrations of added methanol. Nucleic acid, on the other hand, was not detected in the final holdfast fraction despite the lack of a nuclease treatment in the isolation process. This suggests that, despite the holdfast’s reactivity toward nucleic acid dyes and mild sensitivity to DNAse, nucleic acids likely represent a loosely associated component that is lost easily during the extraction process (29, 49).

Imaging of isolated holdfast after staining with fWGA showed that the matrix retained some degree of structural integrity despite the extensive enrichment process. This persistent insolubility points to a mechanism for promoting association between the polysaccharide strands. AFM studies of surface attached holdfasts came to a similar conclusion (28). Two holdfast biosynthesis genes, *hfsH* and *hfsK*, are predicted to encode a polysaccharide N-deacetylase and an N-acetyltransferase respectively, and deleting either gene disrupts the cohesiveness of the holdfast (50, 51). Both enzymes would be predicted to target amino groups from the 4-linked GlcNAc residues detected in our samples, suggesting that a GlcNAc modification pathway increases the structural integrity of the holdfast. Such modifications would have been masked by the acid hydrolysis and acetylation sequence used in the analyses described here. The possibility of GlcNAc modifications represents one of a number of structural questions raised by our results that will require more targeted analytical approaches to address. As a parallel approach, our results should also facilitate *in vitro* characterization of holdfast synthesis enzymes by defining the substrate pool needed for holdfast production.

### Experimental procedures

#### Bacterial strains, growth conditions and genetic manipulations

The strains used in this study are shown in Table 1. For genetic manipulations, *C. crescentus* was grown in peptone-yeast extract (PYE) medium supplemented with 3% (w/v) sucrose, 25µg/mL kanamycin (solid medium) or 5µg/mL kanamycin (liquid medium) when necessary. The plasmid for *hfaB* deletion (pDH108) was created by inserting a fusion with 500bp upstream of the *CC_2629* open reading frame, the first and last twelve nucleotides of *CC_2629* open reading frame, and 500bp downstream of the *CC_2629* open reading frame into pNPTS138 using Gibson assembly. Deletion plasmids for *hfiA* and *hfsJ* have been described previously (32). Unmarked, in-frame deletions were created using a standard two-step procedure with *sacB*-based counter-selection. Cultures for holdfast extraction were prepared by inoculating either M2 salts containing 0.15% (w/v) xylose or PYE with an overnight starter culture of the appropriate strain grown in PYE.

#### Wheat germ agglutinin (WGA) blotting

2.75µL of each sample to be analyzed, along with two-twofold serial dilutions prepared in TU buffer (15mM Tris-HCl pH 8.8 and 4M urea), was spotted onto a nitrocellulose membrane. The membrane was allowed to dry for 30 m, after which it was blocked for a minimum of 1 h using TBST containing 5% (w/v) BSA. The membrane was then stained for a minimum of 1 h using 1.5µg/mL Alexa594-WGA (ThermoFisher) dissolved in TBST containing 1.25% BSA. After washing three times with TBST the membrane was imaged using a BioRad Gel Doc imager on the Alexa647 setting.

#### Phenol extraction of shed holdfast

Δ*hfiA* Δ*hfaB* cells were grown for 24 h in 1 L of M2X or PYE medium. The cell material was removed by centrifuging the culture for 1 h at 17,000 x g. 750 mL of the cell-free spent medium was brought to a concentration of 60% (v/v) ethanol and incubated overnight at 4 °C. The resulting precipitate was isolated by centrifuging for 1 h at 17,000 x g, re-suspended in 110 mL of TES Buffer (10mM Tris-HCl pH 7.4, 2% SDS and 1mM EDTA), and sonicated for 30 s. An equal volume of aqueous phenol was added, and the solution was incubated at 68°C for 15 m. After allowing the phases to separate, the upper aqueous phase was removed and discarded. 80 mL of dH_2_O was added to the remaining phenol phase, the solution was again incubated at 68°C for 15 m and the aqueous phase was again discarded. 12 mL of TEG buffer (10mM Tris-HCl pH 7.4, 4M guanidinium-HCl and 1mM EDTA) was added to the remaining phenol phase. The upper phase was discarded, and the remaining phenol phase was washed a final time with 6 mL dH_2_O. An equal volume of methanol was added to the phenol phase, and the resulting solution was incubated for 1 h at room temperature. The 50% methanol solution was centrifuged for 40 m at 3,000 x g, and the insoluble material was discarded. The supernatant was brought to a concentration of 80% methanol and incubated overnight at −20°C. The 80% methanol solution was then centrifuged at 17,000 x g for 45 m. The pellet was washed with ice-cold 70% ethanol and re-centrifuged, leaving a holdfast enriched pellet.

#### Proteinase K digest

The precipitated phenol phase (above) was resuspended in 5 mL of TU buffer and sonicated for 30 s. Proteinase K was added to 50 µg/mL. The digest was incubated for 2 h at 37°C, followed by a second 2 h incubation at 60°C.

#### Ion exchange chromatography

Proteinase K digested material was loaded on a 1.5 mL Fast Flow Q column that had been equilibrated in TU buffer. The column was then washed with 5 mL of TU buffer and was eluted sequentially with a step gradient of 15mM Tris-HCl pH 8.5 containing 30mM NaCl, 30mM guanidinium-HCl, 300mM guanidinium-HCl and 4M guanidinium-HCl in 3.5 mL fractions. The 300mM GuHCl fraction, which contained the bulk of the WGA reactive signal, was added to 3.5 kDa cutoff dialysis tubing and dialyzed four times against dH_2_O over 24 h. The dialyzed sample was added to a glass tube, frozen at −80°C and lyophilized to dryness.

#### Fluorescence microscopy

Solutions of 10µg/mL AlexaFluor594-WGA and 0.005% Fluoresbrite 0.1µm YG Microspheres were prepared in PYE and used to stain holdfast extracts. 0.5µL of fWGA solution and 1µL of fluorescent bead solution (or 1µL PYE blank) was added to 2.5µL of H_2_O dialyzed holdfast extract. Imaging was performed using a Leica DM500 microscope equipped with a HCX PL APO 63X/ 1.4na Ph3 objective. fWGA staining was visualized using an RFP fluorescence filter (Chroma set 41043) and the fluorescent beads were visualized with a GFP fluorescence filter (Chroma set 41017).

#### Monosaccharide composition analysis

Holdfast samples were hydrolyzed using 2M TFA (200 µL, 2h, 121°C), reduced with NaBD_4_ (10 mg/mL in 1M NH_4_OH, room temperature, overnight), and acetylated using acetic anhydride/pyridine (250 µL + 250 µL, 1h, 100°C). The resulting alditol acetates (AAs) were analyzed on an Agilent 7890A GC interfaced to a 5975C MSD (mass selective detector, electron impact ionization mode). Separation was performed on a 30 m Supelco SP-2331 bonded phase fused silica capillary column (*neutral sugars*) and on a Supelco Equity-1 fused silica capillary column (30 m x 0.25 mm ID) (*amino sugars*). Samples were analyzed both at 100:1 split and 10:1 split in SP-2331, and splitless in Equity-1.

#### Linkage analysis of partially methylated alditol acetates

Holdfast samples were pre-acetylated, permethylated, hydrolyzed, reduced, and re-acetylated. The resultant partially methylated alditol acetates (PMAAs) were analyzed by gas chromatography-mass spectrometry (GC-MS). Before permethylation, the samples were acetylated with acetic anhydride/pyridine (250 µL + 250 µL, 1h, 100°C). Permethylation was performed by two rounds of treatment with sodium hydroxide (15 min) and methyl iodide or deuteromethyl iodide (45 min). Subsequently, the permethylated material was hydrolyzed with 2 M TFA (400 µL, 2h, 121°C), reduced with NaBD_4_ (10 mg/mL in 1M NH_4_OH, room temperature, overnight), and acetylated using acetic anhydride/pyridine (250 µL + 250 µL, 1h, 100°C). The resulting PMAAs were analyzed on an Agilent 7890A GC interfaced to a 5975C MSD (mass selective detector, electron impact ionization mode); separation was performed on a 30 m Supelco SP-2331 bonded phase fused silica capillary column (*neutral sugars*) and on a Supelco Equity-1 fused silica capillary column (30 m x 0.25 mm ID) (*amino sugars*).

## Acknowledgements

We thank members of the Crosson lab for helpful discussions. This work was supported by NIH grant R01GM087353 to S.C and by the Chemical Sciences, Geosciences and Biosciences Division, Office of Basic Energy Sciences, U.S. Department of Energy grant (DE-SC0015662) to Parastoo Azadi at the Complex Carbohydrate Research Center. D.M.H. is supported by the Helen Hay Whitney Foundation.

